# Chimeric origin of eukaryotes from Asgard archaea and ancestral giant viruses

**DOI:** 10.1101/2024.04.22.590592

**Authors:** Sangita Karki, Zachary K. Barth, Frank O. Aylward

## Abstract

The details surrounding the evolution of complex cells remain some of the most enduring mysteries in biology. Recent evidence has demonstrated that Asgard archaea are the closest cellular relatives of eukaryotes, but several eukaryotic enzymes involved in key cellular processes lack phylogenetic affinity with archaea. In particular, phylogenies of eukaryotic DNA and RNA polymerases often support a 3-domain topology that is not consistent with an archaeal origin. Here we present comprehensive phylogenetic analysis of eukaryotic family B DNA polymerases and multimeric RNA polymerases and show that these core subunits of these enzymes are derived from the ancestors of modern giant viruses (phylum *Nucleocytoviricota*). Specifically, we show that the eukaryotic delta polymerase (Polδ), a key processive polymerase required for genome replication in all eukaryotes, clusters within an ancient viral clade, strongly supporting a viral origin. By contrast, the other eukaryotic processive polymerase (Polε), clusters within an Asgard archaeal clade. Together, these observations provide a strong and direct link between early eukaryotes, Asgard archaea, and giant viruses. Lastly, we provide a comprehensive phylogenetic analysis of eukaryotic multimeric RNA polymerases to confirm that RNA polymerase II, which is responsible for mRNA transcription in eukaryotes, is also derived from the ancestors of modern giant viruses. In total, our results support a model of eukaryogenesis in which complex cells emerged from a genomic chimera of Asgard archaea and an ancient viral lineage.

## Main Text

The origin of eukaryotes remains one of the most compelling mysteries in biology. Eukaryotic cells are considerably more complex than those of bacteria and archaea due to the presence of numerous traits including mitochondria, extensive cytoskeletal networks, and an inner membrane system that includes the nucleus and enables the compartmentalization of transcription and translation. Importantly, no eukaryotic organism representing an intermediate level of cellular complexity has been discovered, leading to lively debate regarding the order, timing, and mechanisms through which these complex cellular structures evolved ^1^. In the last decade, the discovery of the Asgard archaea and subsequent evolutionary studies of this group have provided important details regarding the early evolution of eukaryotes ^2,3^. Phylogenetic studies of many core genes involved in translation, ribosome biogenesis, and other key cellular processes often place eukaryotes within the Asgard lineage, and these archaea also encode a range of “eukaryotic signature proteins” that were once thought to be unique to complex cells^3^. Intriguingly, some Asgard archaea also have more complex and dynamic endomembrane systems compared to most other archaea, further implicating them as the likely cellular progenitor to eukaryotes ^4,5^.

Despite these key links between Asgard archaea and eukaryotes, phylogenetic analysis of eukaryotic housekeeping genes sometimes reveals conflicting or weakly-supported links to archaea ^6–11^. This is especially true for eukaryotic genes involved in DNA replication and transcription, which are expected to be core genes present in the last eukaryotic common ancestor^12^. Moreover, the eukaryotic replisome is markedly distinct from that of archaea, which runs counter to expectations if eukaryotes represent direct cellular descendents of Asgard archaea. Due to these discordant phylogenetic signals, it is particularly important to examine the early evolutionary events that led to the emergence of eukaryotic replication and transcriptional machinery, which likely hold clues to how complex cells emerged. Methods for deep-time phylogenetic reconstruction have advanced dramatically in recent years, and the breadth of available genomes to use for evolutionary analyses has dramatically expanded, creating an opportune time to evaluate the origin of these key eukaryotic complexes.

Importantly, molecular studies have shed light on the details of processive DNA replication in eukaryotes, and it is now clear which enzymes play central roles in whole-genome synthesis ^13^. Eukaryotic genome replication is a complex process that is performed by a suite of DNA polymerases and accessory factors, and close examination of eukaryotic replisome evolution with special emphasis on processive polymerases is therefore critical to evaluate the process through which whole-genome replication evolved in complex cells. Eukaryotes encode four family B DNA polymerases (PolBs), and molecular studies have shown that Polε and Polδ perform the majority of leading and lagging strand synthesis ^8,13^. The two other PolBs – Polα and Polζ – have major roles in replication initiation and DNA repair, respectively. The DNA sliding clamp–also called the Proliferating Cell Nuclear Antigen, or PCNA–is a key component of the replisome that associates with both delta and epsilon polymerases and prevents them from dissociating from DNA during polymerization, effectively providing the processivity that is needed for replication of large cellular genomes ^13^.

To shed light on the evolutionary origins of the eukaryotic replisome components, we performed comprehensive phylogenetic analysis of both cellular and viral PolBs with the primary goal of elucidating the evolutionary origins of Polδ and Polε (Figure 1A; see Methods). We included as broad a sampling of PolB enzymes as possible in order to provide an accurate reconstruction of ancient evolutionary events. Importantly, we included PolBs from eukaryotes and archaea, as well as several distinct lineages of large DNA viruses, such as herpesviruses (order *Herpesvirales*), giant viruses (phylum *Nucleocytoviricota*), and members of the recently-discovered phylum *Mirusviricota* ^14^. For multi-sequence alignment we used the newly-developed Muscle5 algorithm, which has been shown to substantially improve the alignment of divergent proteins ^15^. Moreover, to ensure that the topology of our trees was well-supported, we employed both regular and complex substitution models in our phylogenetic analyses (LG+F+R10 and LG+C60+F+R10, respectively), we employed different levels of taxon sampling (Figure 1 A and B and Extended Data Figure 3A), and we performed phylogenetic reconstruction on alignments that were trimmed to several different levels of stringency (Extended Data Figure 7). In all of our resulting trees we found that Polδ formed a distinct clade sister to the *Nucleocytoviricota* and nested within a broader clade that includes the herpesviruses and *Mirusviricota* (Figure 1). This result was well-supported in all of the trees that we constructed (>99% ultrafast bootstrap support in all cases).

**Figure 1:**
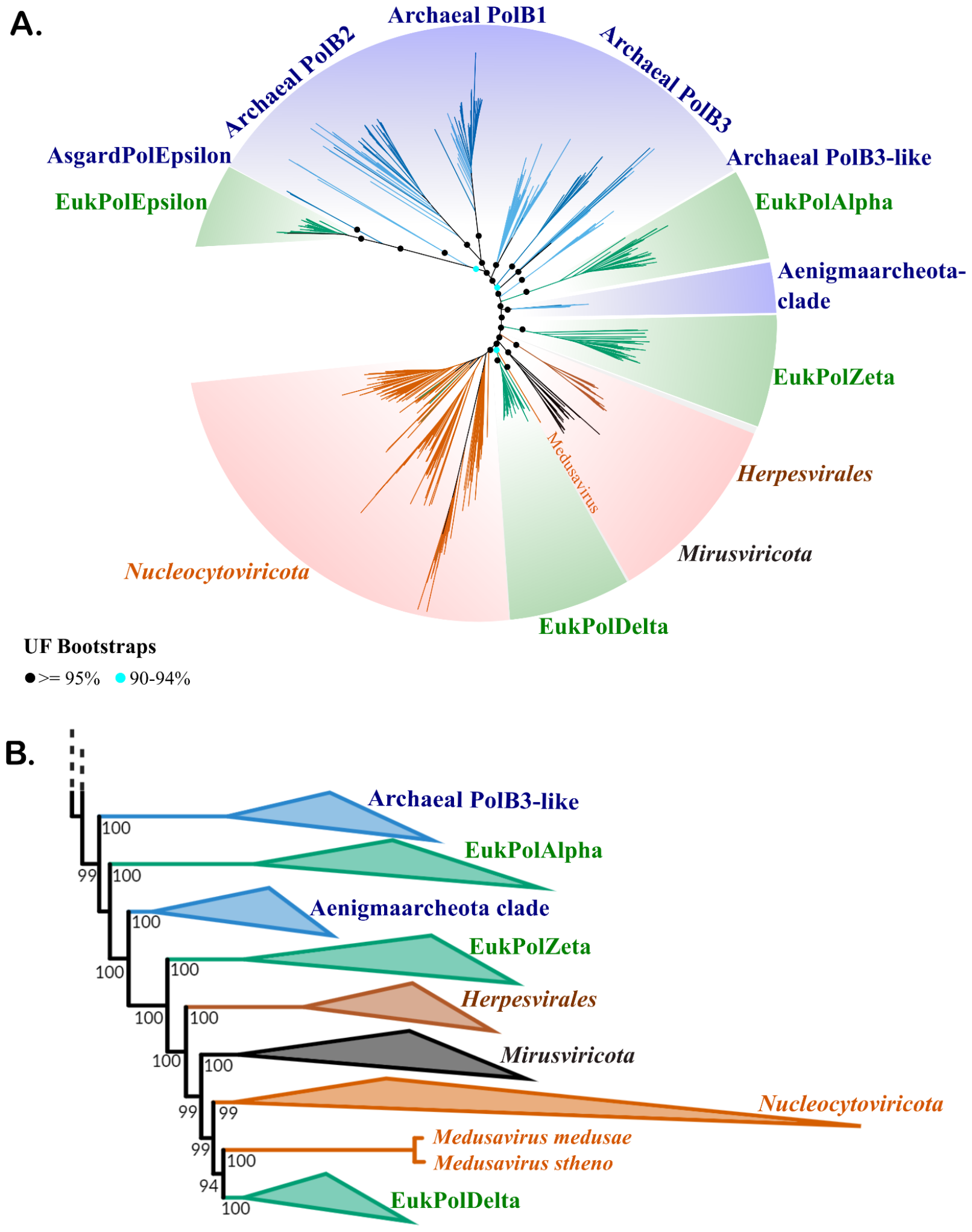
Phylogenetic tree of DNA polymerase family B demonstrating nested placement of Polδ in a viral clade and Polε with Asgard archaea. (957 total sequences, 1417 sites). A) Maximum-likelihood analysis was performed using IQ-TREE using the complex model LG+C60+F+G. Ultrafast bootstrap support values for select deep-branching nodes are shown (black dot >=95%, blue dot 90-94%). For clarity, support values are only provided for select internal nodes. B) Rectangular representation of the region of the polB phylogenetic tree highlighting the evolutionary relationships between viral groups and eukaryotic Polδ. Values at nodes represent their ultrafast bootstrap support.

Fast-evolving sites may introduce noise and obscure phylogenetic inference in protein families ^16^, and so to further confirm the topology we made multiple sets of PolB trees in which increasing numbers of fast-evolving sites were iteratively removed from the alignment (see Methods for details). The overall topology of these trees remained consistent until 40% of all alignment positions were removed, after which the overall quality of the tree deteriorated as evidenced by the collapse of monophyly in major clades of archaea and viruses (Extended Data Figure 4A). This analysis provides another confirmation of the nested placement of Polδ within a viral clade and demonstrates that this topology is not an artifact of fast-evolving sites in the alignment. Importantly, this finding is consistent with earlier studies that examined a smaller set of PolB proteins and provided the first insight that Polδ may be viral-derived ^17^. Moreover, our results also show that viruses of the genus *Medusavirus* have basal placement to the Polδ clade, consistent with previous phylogenetic analysis ^18^ and clearly supporting a nested placement of eukaryotic proteins within a viral clade that would be expected under a scenario of viral origin.

By contrast, the other processive PolB in eukaryotes, Polε, formed a clade with Asgard archaea, also with high support in all cases (100%) (Figure 1, Extended Data Figures 2A, 3A, 4A, and 7). The Asgard archaeal ε-like polymerases were encoded primarily by members of the Hodarchaeota and Heimdallarchaeota (Supplementary Table 2). The combined placement of Polδ and Polε provides a direct link between Asgard archaea and the ancestor of giant viruses in the early evolution of eukaryotes. Together, these results support a chimeric origin of processive DNA replication in eukaryotes that arose from a combination of enzymes found in both the ancestor of modern giant viruses and Asgard archaea.

The sliding clamp (PCNA) associates with both polymerases δ and ε during DNA replication and is a key component of processive replication that is needed for whole-genome synthesis. Due to the key role of the DNA sliding clamp in replication processivity, we also performed exhaustive phylogenetic analyses on viral and cellular homologs of this protein, using methods similar to those that we employed for the PolB phylogenies (see Methods). Our results also suggest a viral origin of the eukaryotic sliding clamp (Extended Data Figure 1), but these results are more tentative owing to the relatively weak phylogenetic signal in this protein compared to PolB (mean length of ∼270 aa for PCNA compared to >1000 aa for most PolBs). Nonetheless, our analysis recovered placement of eukaryotic PCNA nested within a broader viral clade and distal to archaeal homologs.

To examine the evolutionary origins of transcription in eukaryotes, we then sought to broaden our analysis to include multimeric RNA polymerase (RNAP). RNAP is a key enzyme in which the two major subunits are found in a single copy in bacteria, archaea, and some DNA viruses, and three copies in eukaryotes (referred to as I, II, and III). Importantly, a previous study of cellular RNAP found only weak or ambiguous support for an archaeal origin ^6^, and another study has proposed a viral origin of eukaryotic RNAP II and RNAP III ^9^. These phylogenetic signals that contrast with an Asgard archaeal origin are notable considering that the two largest RNA polymerase subunits are large proteins (∼1500 aa) that are expected to provide one of the strongest phylogenetic signals amongst core protein sequences ^19^. We therefore examined the evolutionary origin of this enzyme in detail using an updated genomic representation of both viral and cellular proteins, including those from the recently-discovered phylum *Mirusviricota*. Using a complex substitution model, we confirmed that eukaryotic RNAP II is nested within a viral clade (LG+C60+F+G; 100% bootstrap support), but that RNAP I and III form deep-branching groups with unclear origin (Figure 2). Similar to our PolB analyses, we confirmed this result with extensive testing of alternative phylogenetic models, alignment trimming severity, and taxon sampling (see Methods). It is notable that RNAP II is responsible for mRNA transcription in eukaryotes, which is essentially the same role that this enzyme plays in giant viruses. This strongly suggests that, similar to Polδ, RNAP II was acquired from an ancient viral ancestor of modern giant viruses.

**Figure 2:**
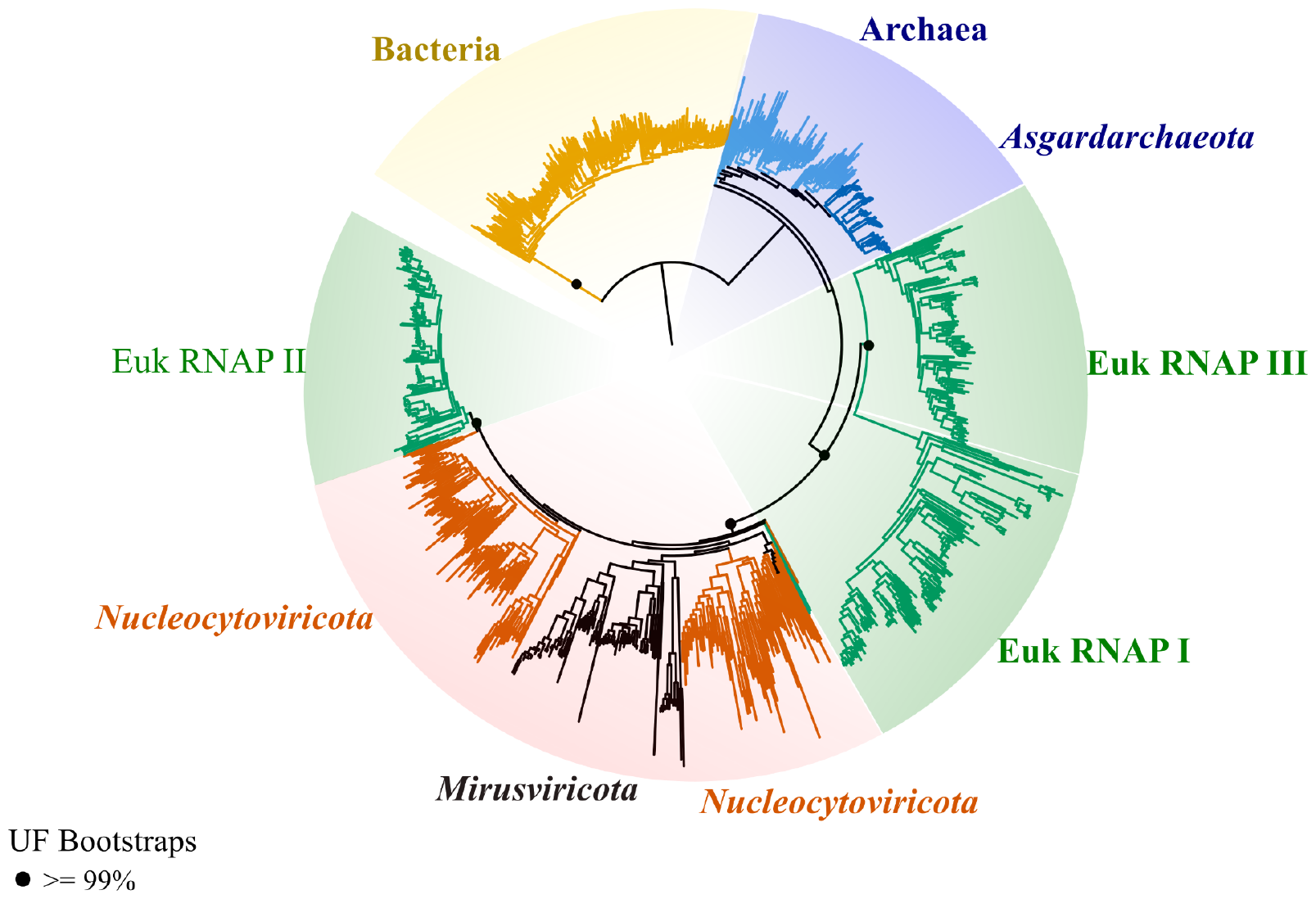
Phylogenetic tree for RNA Polymerase (RNAP) showing placement of eukaryotic RNAP nested within a viral clade. The alignment is based on a concatenated set of Beta and Beta prime subunits from 1017 sequences (resulting in a total alignment length of 3812 sites). Maximum-likelihood analysis was performed using IQ-TREE under a complex model (LG+C60+F+G). The dots on the branches represent ultrafast bootstrap support values (black dot >= 99%). For clarity, support values are only provided for select internal nodes. Full trees are available in the supplemental material. The tree is rooted between the bacteria and all other taxa.

The evolutionary origins of RNAP I and RNAP III, which are involved primarily in rRNA and tRNA transcription, remain unclear in our phylogenetic analysis. It is notable that the branches leading to the diversification of RNAP I, RNAP III, and viral/eukaryotic RNAP II are extremely short, indicative of a rapid evolutionary transition that occurred at or around the time of eukaryogenesis. We postulate that eukaryotic RNAP I and III emerged through a process of duplication from the ancestral viral-derived RNAP II, and that phylogenetic placement of these enzymes is complicated by the rapid evolution and neofunctionalization that took place shortly after duplication. To potentially resolve this branching order, we constructed a series of RNAP trees in which a range of fast-evolving sites were removed, but we did not see any substantial change in topology (Extended Data Figure 4B). Resolving the branching order of ancient evolutionary events that occurred close together in time is notoriously difficult ^20^, and we predict that assessing the relative times at which RNAP I, RNAP II, and RNAP III emerged will remain an important challenge for years to come.

The theory that viruses have played a role in eukaryogenesis goes back decades, but it has remained difficult to test owing to the challenges of deep phylogenetic inference coupled with the lack of sufficient viral sequences to include in evolutionary analyses. Most early theories of viral eukaryogenesis focused on a possible viral origin of nuclear processes such as replication, transcription, and mRNA processing ^21–23^, stemmed in part from early phylogenetic analysis of family B DNA polymerases that suggested a possible phylogenetic affinity between eukaryotic enzymes and some distant viral homologs ^21,23^. Subsequent studies lent further support to a link between Polδ and viral polymerases ^17,18^, and yet it has remained challenging to confidently resolve these evolutionary relationships and confirm the direction of these ancient gene transfer events. These challenges arose in part because few viral genomes were available and only a small number of viral polymerases could be included in phylogenetic analyses, but the dramatic expansion of the known diversity of giant viruses ^24,25^ coupled with the discovery of the phylum *Mirusviricota* ^*14*^ have created an unprecedented opportunity to examine the deep evolutionary roots of eukaryotic enzymes in the context of this broader viral diversity. Importantly, *Mirusviricota* form a distinct branch that is basal-branching to eukaryotes in our PolB tree, and inclusion of these sequences is therefore key to establishing the placement of eukaryotic Polδ within a well-defined viral lineage.

With the exception of Polε, our results show that the genes required for processive DNA replication and transcription are primarily viral-derived (Figure 3A), while other key processes in eukaryotes such as translation, the ubiquitin-proteasome system, and endomembrane trafficking have been shown to have primarily archaeal origins ^2,3^ (Figure 3A). Although it is challenging to disentangle the exact scenario of gene transfer that led to this pattern, it is striking to note that our results are remarkably consistent with a model of viral eukaryogenesis previously put forward by Bell ^26^. In this model, the eukaryotic nucleus is derived from a membrane enclosed “virus factory” (VF) that formed during a chronic infection (Figure 3B). In giant viruses, the enzymes involved in replication and transcription are encoded by the virus and typically occur in virus factories that form during infection ^27^, while extra-nuclear functions are primarily provided by the host cell. In Bell’s model, an Asgard archaea is infected by a large DNA virus ancestral to modern giant viruses, leading to formation of a VF within a compartment derived from the cellular endomembrane system. Transfer of essential functions from the archaeal genome to the VF would likely be one of the first steps in the domestication of the VF, and it would link viability of the cell to retention of the proto-nucleus. Nuclear import of genes from mitochondria and chloroplasts has been well-documented across eukaryotes, providing important precedent for this model of nuclear import and domestication.

**Figure 3.**
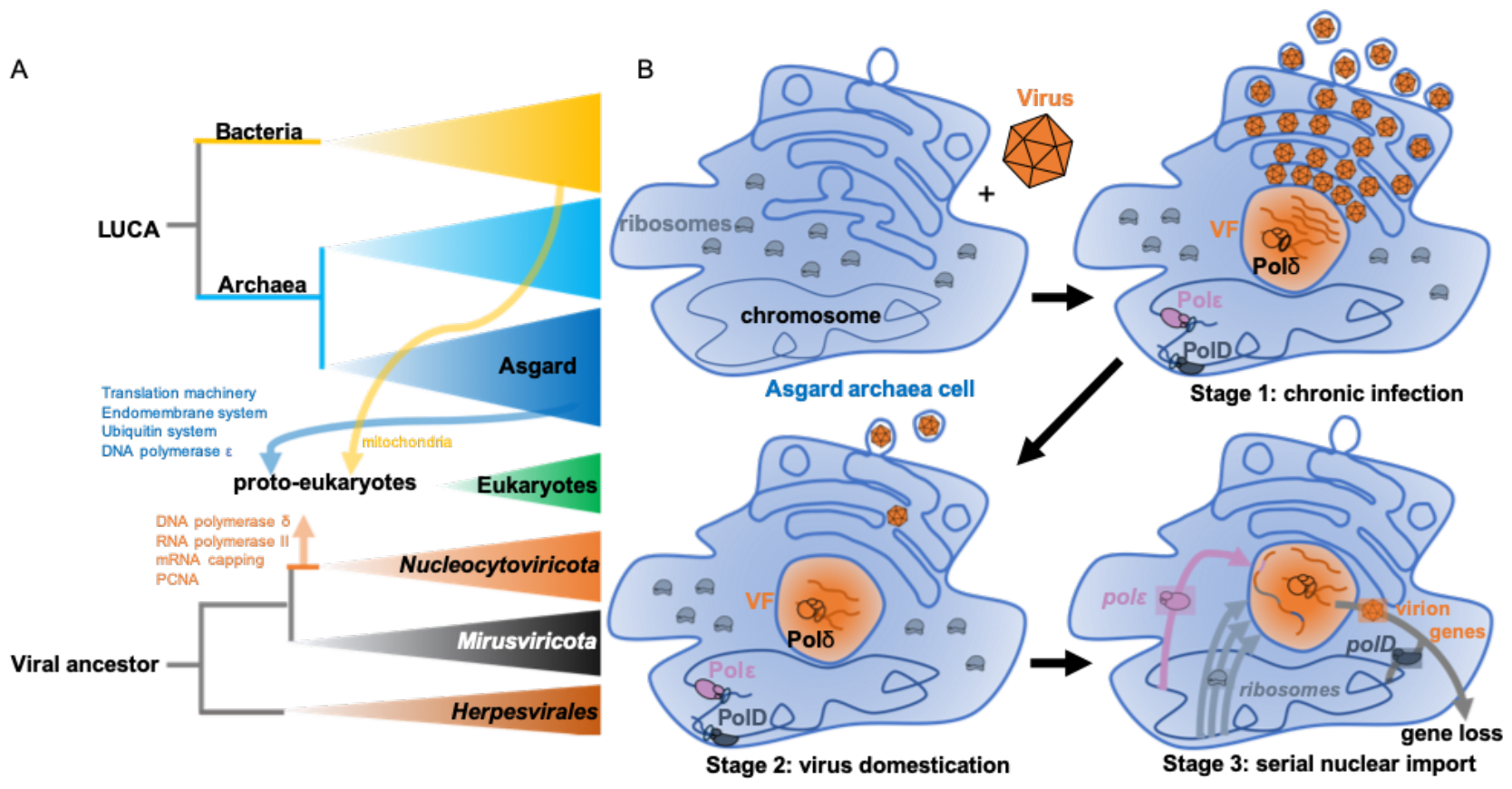
Chimeric phylogenetic signals in eukaryotes derive from both Asgard archaea and the ancestors of giant viruses. A) Diagram showing the genetic contributions of separate lineages based on our results as well as current literature. B) A model of viral eukaryogenesis that is consistent with the phylogenetic evidence. (Top left), The first eukaryotic common ancestor cell is a Heimdallarchaeon cell with a complex endomembrane system. Stage 1 of eukaryogenesis is a chronic infection by a virus that forms a viral factory (VF) from the cellular endomembrane system. Viral DNA within the VF is spatially separated from the ribosomes within the host cytosol. The ancestor of eukaryotic Polδ facilitates viral DNA replication within the VF, while the cellular chromosome is maintained by Polε and archaeal PolD. Stage 2 consists of viral domestication. The virus evolves to favor vertical inheritance over horizontal propagation, while being maintained as an episome within the VF. Stage 3 is serial nuclear import. Essential genes are horizontally transferred from the cellular chromosome to the compartmentalized viral genome, making the virus an essential component of the cell. Over many generations, the ribosomal genes and eukaryotic signature proteins derived from the cellular ancestor are all imported into the viral factory, while signature proteins of Asgard archaea, such as PolD, and the virion structural genes are lost.

The import of ribosomal genes to the nucleus would be a critical step in VF domestication, while the import of genes for transcription and replication would be less important owing to their presence in the viral-derived nuclear genome already. The archaeal origin of Polε suggests that this enzyme was also imported into the proto-nucleus and became part of the replisome together with Polδ during this process. Consistent with the model of viral eukaryogenesis, several lines of evidence suggest that Polε was likely a later addition to the eukaryotic replisome, chief among them that Polε’s catalytic activity is non-essential for DNA synthesis ^28–30^ and that it does not require PCNA for processivity as Polδ does ^13^ (see supplementary discussion). PolD, the primary processive polymerase in most archaea, was presumably not imported into the nucleus and eventually lost because its function was redundant with the ancestral Polδ homolog already in the nucleus. Another key step in eukaryogenesis would have been the acquisition of the mitochondria, which according to this model would have occurred later after the origin of the nucleus. This is consistent with previous phylogenomic evidence for a late mitochondrial acquisition that occurred in a host that already had chimeric ancestry ^31^.

This work is the latest in a string of recent discoveries that have lent greater plausibility to the viral eukaryogenesis hypothesis. Historically, a shortcoming of viral eukaryogenesis has been that the VF of extant giant viruses is derived from the endoplasmic reticulum of the host cell, and complex endomembrane systems were unknown outside of eukaryotes. Now, complex endomembrane systems have been observed in multiple lineages of archaea ^4,5^, suggesting that complex endomembrane systems existed prior to the evolution of a nucleus and could be co-opted to form VFs in archaeal cells. Secondly, viral compartmentalization of DNA replication and transcription has also evolved in viruses that infect bacteria ^32^, demonstrating that nucleus-like structures form in associations between viruses and prokaryotes. Finally, archaea are also known to host chronic infections where infected cells are able to produce both free virions by budding as well as new infected and uninfected cells by asymmetric cell division ^33^, establishing that some archaeal viruses have the capacity for long-term non-lytic associations with cells. Indeed, although chronic infection has often been understudied owing to technical difficulties in characterizing these associations, it is likely a common strategy employed by numerous viruses of bacteria and archaea^34^.

The viral origin of key components of the eukaryotic replisome and transcriptional apparatus present a potential resolution to the paradox of discordant phylogenetic signals in eukaryotic genomes. Viral eukaryogenesis also offers an explanation for how eukaryotes evolved a replisome so markedly distinct from archaea. Most current models of eukaryogenesis place Asgard archaea as the closest living relatives of modern eukaryotes ^2,3^, but it has remained mysterious why genes involved in transcription and DNA replication have complex evolutionary histories that conflict with this model ^6–8^. Our results provide strong support for a viral origin for key components of eukaryotic replication and mRNA transcriptional machinery, which, together with an Asgard archaeal origin for the ribosome and other eukaryotic signature proteins, provides a potential resolution to this conundrum. These results also bring the importance of understanding viral diversity in the biosphere into sharp focus. Indeed, as additional lineages of eukaryotic and archaeal viruses are discovered, we predict that further clues regarding the early evolution of eukaryotes will be revealed. In particular, the discovery of large archaeal viruses that encode their own transcription and replication machinery or form nucleus-like structures during infection would lend further support to the model of viral eukaryogenesis described here.

## Methods

### Dataset Compilation

We compiled a set of high-quality bacterial, archeal, and eukaryotic and viral genomes for subsequent phylogenetic analyses. For eukaryotic genomes, we used all genomes available on the eggNOG v5.0 database ^35^. To increase the representation of unicellular eukaryotes, we also included seven complete or chromosome-level genomes of protists available on the National Center for Biotechnology Information (NCBI) databases ^36^ as of October 8, 2021. For bacterial and archaeal genomes, we retrieved genomes from the Genome Taxonomy Database (GTDB, v95)^37^. To enrich our database in Asgard archaeal genomes, we also included genomes from this group that were reported in a recent large-scale comparative genomic study ^38^ that were not already present in the GTDB. For viral lineages, we focused on members of the *Herpesvirales, Nucleocytoviricota* (i.e. “giant viruses”), and the recently-discovered phylum *Mirusviricota*. We used complete herpesvirus genomes available in NCBI as of July 2023, all nucleocytovirus genomes available in the Giant Virus Database (GVDB) (https://faylward.github.io/GVDB/) ^39^, and all mirusvirus genomes published in a recent study ^14^. For the PolB analysis we also considered including sequences derived from other viral groups that encode this enzyme, such as some tailed phages (class *Caudoviricetes*), adenoviruses, baculoviruses, polinton-like viruses, virophages, and some recently-discovered viruses of Asgard archaea ^40^. In initial diagnostic trees that we constructed for PolB (see methods below) these sequences formed long branches that clustered with archaeal PolB1, 2, and 3 clades, and we therefore removed them from our final analysis on the grounds that these long branches could compromise the overall topology of the tree. Moreover, the PolBs from most of these viral groups are protein-primed (not processive), and therefore not as relevant to our analysis given the focus of our work on processive DNA polymerase evolution. For eukaryotic genomes, we used protein predictions already available on EggNOG v. 5.0, and for all other taxa we predicted proteins using Prodigal v. 2.6.3 with default parameters^41^.

### Sampling of taxa

Highly biased taxon sampling can adversely affect phylogenetic inference ^19^. We therefore sought to balance the number of different lineages used in our subsequent phylogenetic analyses by sub-sampling groups of over-represented lineages, which for our purposes were bacteria, archaea, plant, metazoans, fungi, and giant viruses. For bacteria and archaea, we chose high-quality representative genomes from each class in the GTDB to include using a methodology described previously. For eukaryotes, we manually selected a subset of 127 genomes to include in order to remove the overabundance of genomes from the Fungi, Opisthokonta, and Viridiplantae lineages in the EggNOG database, and we added 7 complete or chromosome-level genomes of protist lineages from the NCBI. For nucleocytoviruses, we down-sampled the full set of 1,381 genomes in the GVDB to 343 by including only genus-level representatives from the taxonomy available in this database. For this down-sampling, we chose the genome of the genus-level representative with the highest N50 contig length. We did not down-sample mirusviruses and herpesviruses because relatively few genomes from these lineages were already available. After this down-sampling, we arrived at a genome set that included 127 eukaryotes, 279 archaea, 230 bacteria, 343 nucleocytoviruses, 111 mirusviruses, and 113 herpesviruses. These genomes were a starting point for phylogenetic inference of all trees in our study, and most of the trees that we subsequently analyzed did not include all of these taxa because some lineages lack certain proteins (e.g. most bacteria do not encode family B DNA polymerases). A full list of all genomes used is available in https://zenodo.org/records/10956246 and Supplementary Table 1.

### Dataset Curation and quality check

For prediction of PolB and PCNA homologs in our genome set, we used a custom python script that uses the hmmsearch command in HMMER3^42^ (see Code Availability section). For Hidden Markov Model (HMM) references we used PolB and PCNA models from Pfam v. 32.0^43^ (accessions PF00136 and PF00705 respectively). For multi-subunit RNA polymerase (RNAP), we used the markerfinder_v2.py script to both identify homologs of the beta and betaprime subunit of RNAP and then concatenate them together into a single alignment. For eukaryotes, we did this by matching to custom HMMs that we designed for these subunits in RNAP I, II, and III. For identification of beta and betaprime RNAP subunits in bacteria, archaea, and viruses, we used the COG0085 and COG0086 HMMs designed previously^19^. For all trees, prior to alignment we first dereplicated nearly-identical sequences using CD-HIT version 4.8.1^44^. For PolB trees, we also removed all sequences <650 aa on the grounds that these were likely truncated or erroneously predicted. For the RNAP tree, we did not include taxa in the analysis unless both the beta and betaprime subunit could be identified and included in the alignment.

### Phylogenetic tree reconstruction and benchmarking

For all alignments we used Muscle5^15^ (parameters “-super5” for input sequences), which has recently been shown to substantially improve multi-sequence alignment compared to previous methods. For RNAP specifically, we used a custom script merge_and_align.py that uses Muscle5 algorithm to align and then concatenate the RNAP protein sequences (see code availability section). We trimmed the alignments with trimAl v1.4. rev15^45^ (parameter -gt 0.1, but see below for alternative trimming strategies). We manually inspected all alignments with AliView^46^ and removed sequences with long, continuous gaps that may hinder phylogenetic inference. In these cases, alignment was then re-performed and the alignments were inspected again. In the case of PolB, we inspected the untrimmed alignments and found some long insertions in some sequences that correspond to inteins, but we confirmed that these were removed by subsequent trimming steps.

For all the gene trees (PolB, RNAP, and PCNA), we initially constructed diagnostic phylogenetic trees using IQ-TREE v1.6.12^47^ with the option -bb 1000 to generate 1,000 ultrafast bootstraps^48^, - m MFP to determine the best-fit model^49^, -nt AUTO and --runs 5 to select the highest likelihood tree. These initial trees were inspected, and long branches that represent rogue taxa or low-quality sequences were removed (<10 sequences from each tree) prior to re-alignment. Moreover, upon inspecting the initial diagnostic trees, we noticed that several large clades of giant viruses and archaea were present, and we randomly down-sampled these clades by 20% using the seqtk subseq function to lessen the computational burden and further prevent biased taxon sampling across the tree. We also noticed that poxviruses had unstable placement in our diagnostic trees, consistent with previous findings ^9^, and we therefore removed this lineage from further analyses. After rogue taxa removal and the last round of subsampling, alignment and trimming procedures were run again.

Once the final alignment was obtained, we then reconstructed maximum likelihood phylogenetic trees using IQ-TREE (parameters -bb 1000, -m MFP, -nt AUTO, –runs 5). The LG+F+R10 model was selected as best fit substitution model based on Bayesian Information Criterion (BIC) for the PolB and RNAP tree, while LG+F+R7 was chosen as best fit for the PCNA tree by ModelFinder (-m MFP). Because amino acid substitution rates likely vary across alignments, we also inferred trees using complex models (C-models) that have different substitution matrices for every position in the alignment^50^ (LG+C60+F+G). We then compared our models from the -MFP option to the most complex C60 model. Although the trees inferred with the -MFP option generally had lower BICs, we still examined the trees inferred with a complex model (LG+C60+F+G) to assess any differences in topology that could be detected using the different methods.

### Further phylogenetic tree validation

We performed several tests to examine how alignment trimming severity, removal of fast-evolving sites, and taxon sampling affected our phylogenetic inference. To examine if different trimming methods could impact the topology of our PolB or RNAP trees, we re-made these trees using more stringent levels of alignment trimming (see Extended Data Figures 6 and 7). Our original trimming strategy was to remove all sites with >90% gaps (-gt 0.1 option in trimAl), and so for more stringent trimming we removed all sites with 50% or more gaps (-gt 0.5 parameter) or by using the automated trimming stringency (-automated1 option). For PolB this resulted in alignment lengths of 867aa (for -gt 0.5) and 351aa (for -automated1) compared to 1417aa for the primary alignment. For our concatenated RNAP alignment this resulted in alignment lengths of 2420aa (for -gt 0.5) and 1117aa (for -automated1) compared to 3812aa for the primary alignment that we used in our analysis. After generating these alternatively trimmed alignments we inferred phylogenies in IQ-TREE using the same LG + F+R10 substitution model as determined by ModelFinder.

Taxon sampling has been shown to impact phylogenetic tree inference, and we therefore sought to examine if the topology of our trees were consistent when using a smaller set of taxa. To test the effect of taxon downsampling, we downsampled the PolB and RNAP protein sequences by ∼50% to 375 sequences for PolB and 517 sequences for RNAP, while keeping the overall proportion of cellular and viral groups consistent (see Extended Data Figure 3). We then generated alignments with Muscle5, used the same alignment QC procedure described for our original trees, and generated trees in IQ-TREE using the LG + F+R10 substitution model.

Lastly, we sought to examine if the removal of fast-evolving sites would alter the topology of our trees (see Extended Data Figure 4). It has been suggested that the removal of fast-evolving sites helps to increase the signal-to-noise ratio in phylogenetic inference ^51^, although a recent study has indicated that fast-evolving sites are informative for tree-building ^52^. We therefore inferred site-specific evolutionary rates from our trimmed PolB and RNAP primary alignments using the -wsr parameter in IQ-TREE v1.6.12^47^. This produced ten different rate categories, which we then sequentially removed before inferring trees with the LG + F+R10 model.

## Code availability

PolB and PCNA homolog identification was done using the custom python script hmmsearch_wrapper: https://github.com/sangitakarki/hmmsearch_wrapper.

RNAP subunits were identified and concatenated alignments were generated using markerfinder-euk: https://github.com/faylward/markerfinder

## Data availability

All genomes and alignments used in this study can be found here: https://zenodo.org/records/10956246. All phylogenetic trees are available on interactive Tree of Life (iTOL): https://itol.embl.de/shared/15ttJikbnoVmi and https://itol.embl.de/shared/1l6saIRHqS5eY

