## Extended Data for "Chimeric origin of eukaryotes from Asgard archaea and ancestral giant viruses"

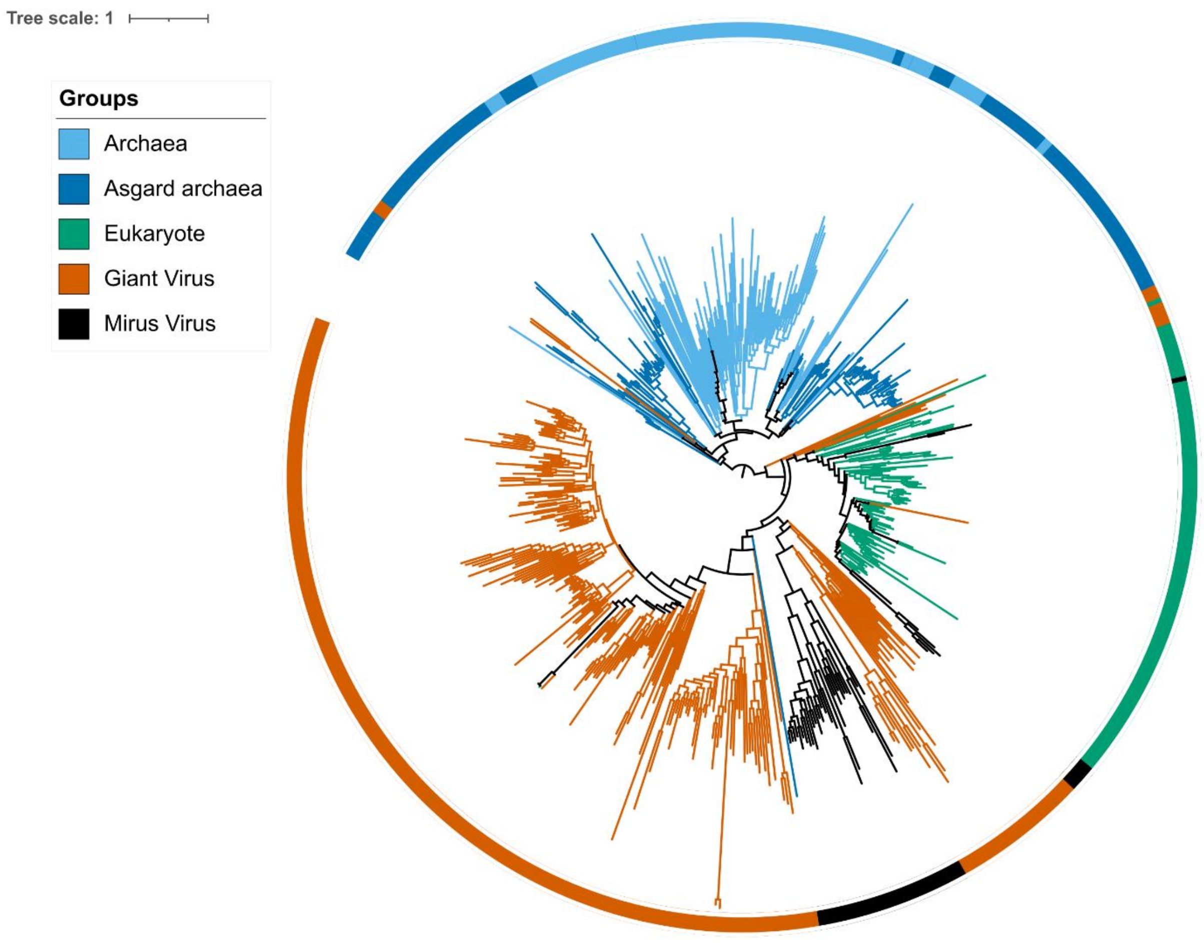


Extended Data Figure 1 | Phylogenetic tree for DNA Sliding Clamp (PCNA) for 606 sequences (resulting in total alignment length of 289 sites). Maximum-likelihood analysis was performed using IQ-TREE under a complex model (LG+C60+F+G). The tree is rooted within an archaeal group. The tree shows eukaryotic sliding clamp clustered within the viral group.

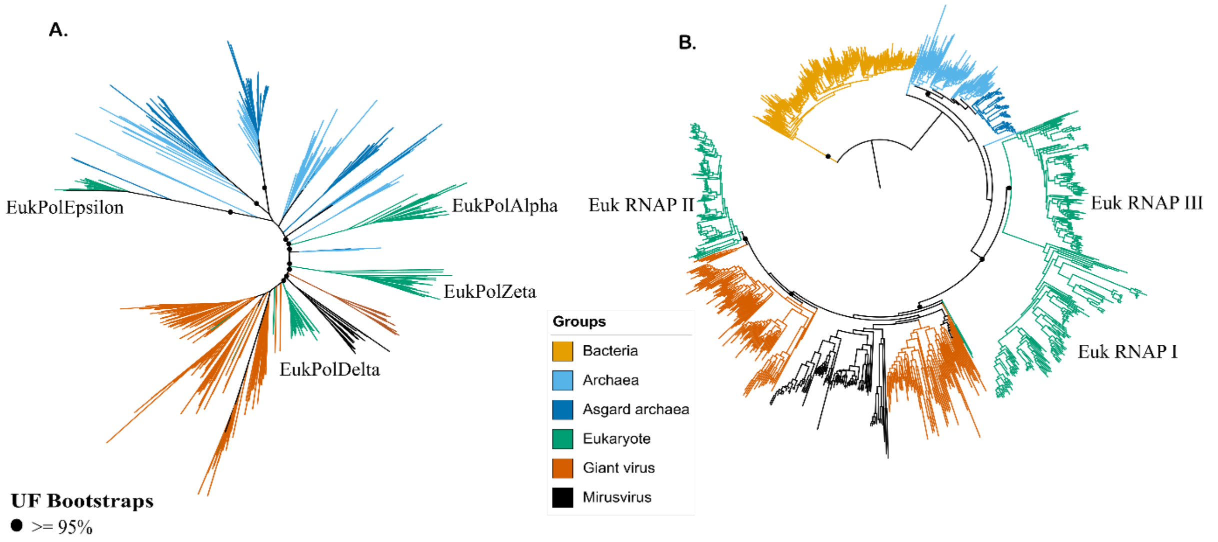


Extended Data Figure 2 | Phylogenetic tree for DNA Polymerase B and RNA Polymerase. The main text figures show trees with complex model while these trees are inferred using regular model. The trees were inferred using the LG+F+R10 model that was chosen as best fit by ModelFinder (-MFP). A. DNA Polymerase B tree is unrooted while B. RNA Polymerase is rooted within bacteria. Dots on the main nodes represent ultrafast bootstrap support.

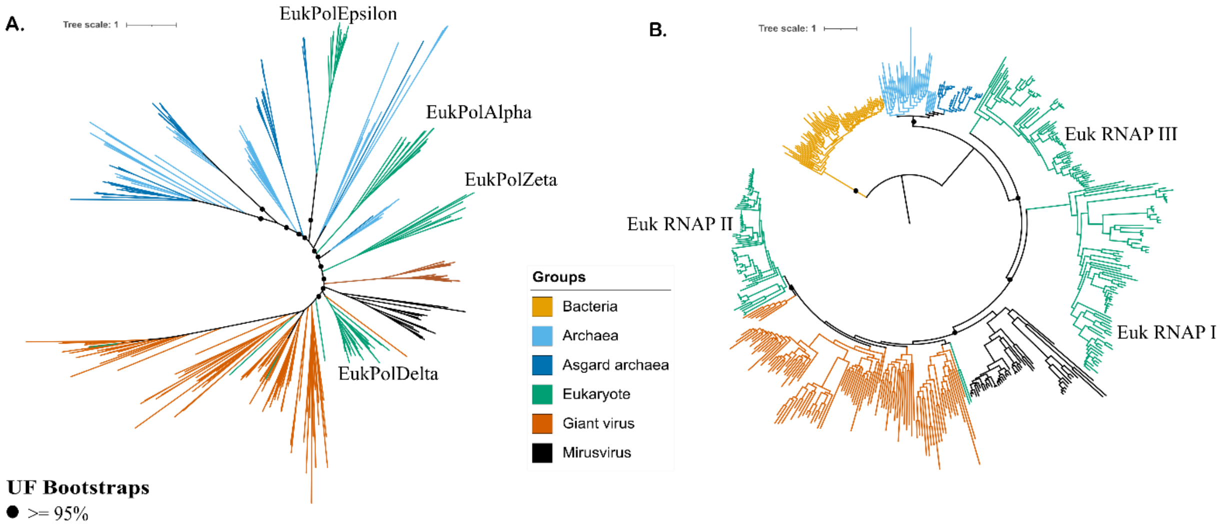


Extended Data Figure 3 | Phylogenetic tree for DNA Polymerase B and RNA Polymerase reduced sets (375 PolB sequences and 517 RNAP sequences vs. 957 PolB sequences and 1017 RNAP sequences previously). The trees were inferred using the LG +F+R10 model. A) DNA Polymerase B tree is unrooted while B) RNA Polymerase is rooted within bacteria. Dots on the main nodes represent ultrafast bootstrap support.

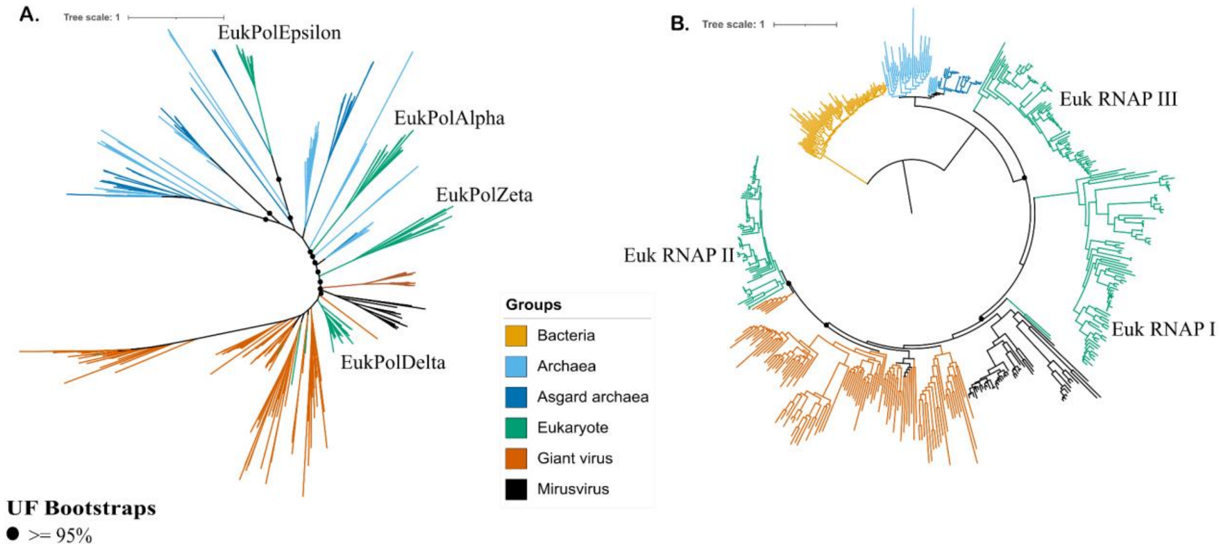


Extended Data Figure 4 | Representative phylogenetic tree after removing 30% of the fast-evolving sites (0-slowest, 10-fastest). The topology is still intact and well supported for viral derives. The trees were inferred using the LG +F+R10 model. A. DNA Polymerase B tree is unrooted while B. RNA Polymerase is rooted within bacteria. Dots on the main nodes represent ultrafast bootstrap support.

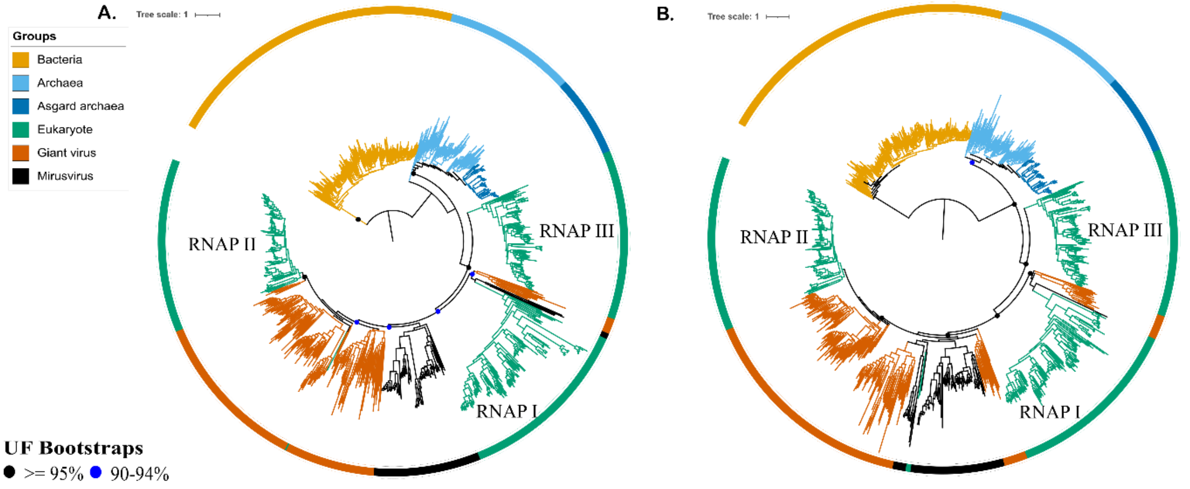


Extended Data Figure 5 | Individual phylogenetic tree for Beta and beta prime subunits of RNA polymerase. The trees were inferred using the LG +F+R10 model. Both trees are rooted within Bacteria. A. RNA polymerase Beta subunit. B. RNA polymerase Beta prime subunit. Dots on the main nodes represent ultrafast bootstrap support.

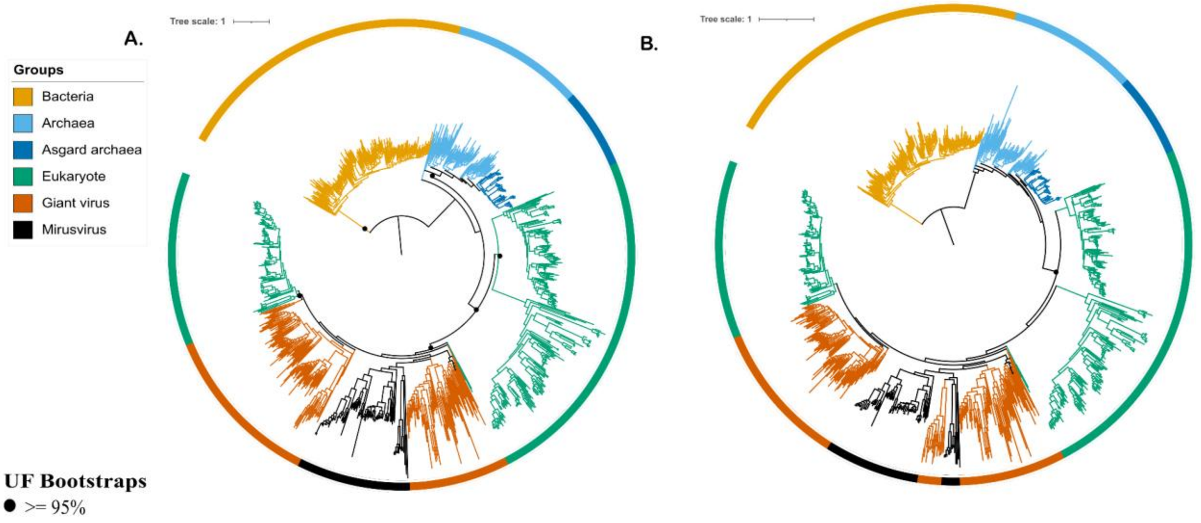


Extended Data Figure 6 | Phylogenetic tree for concatenated RNAP tree using different trimming parameters. A. trimmed parameter -gt 0.5 B. trimmed parameter -gt automated1. Both trees were inferred using the LG +F+R10 model. Both trees are rooted within Bacteria. Dots on the main nodes represent ultrafast bootstrap support.

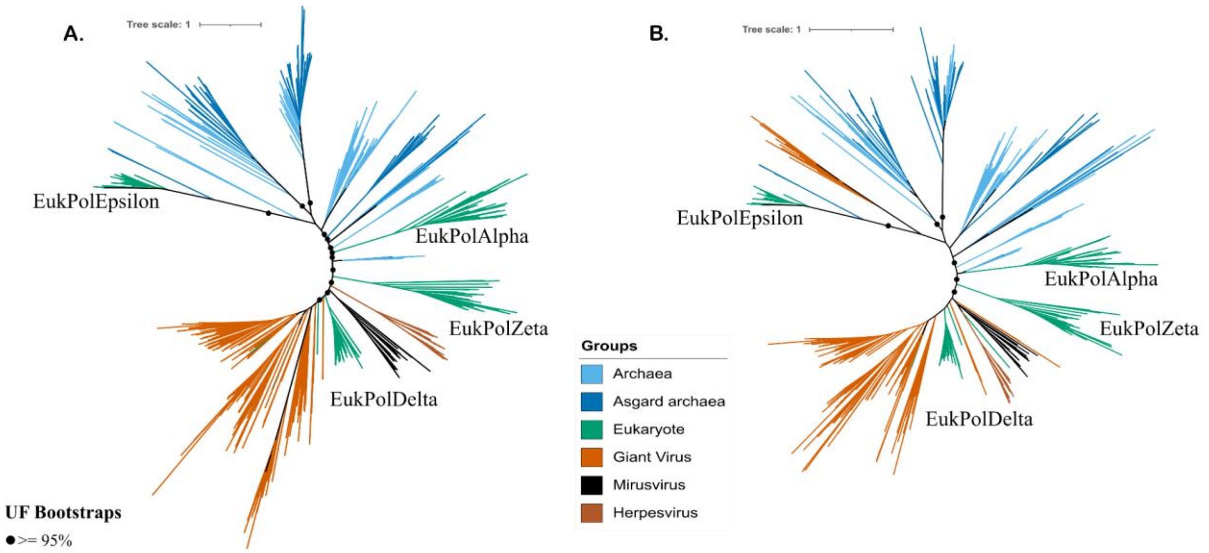


Extended Data Figure 7 | PolB phylogenetic tree using different trimming parameters. A. trimmed parameter -gt 0.5 B. trimmed parameter -gt automated1. Both trees were inferred using the LG +F+R10 model. Dots on the main nodes represent ultrafast bootstrap support.
