## Supplemental Text for "Chimeric origin of eukaryotes from Asgard archaea and ancestral giant viruses"

#### The domain organization and phylogenetic signals of eukaryotic PolEpsilon suggest horizontal acquisition

In eukaryotic DNA replication, there is a division of labor where Polε and Polδ primarily replicate the leading and lagging strand, respectively [1]. Recent work has shown the presence of Polε homologs in some Asgard archaea [2,3], the assumed ancestors of eukaryotic cells. Despite the phylogenetic link between Asgard archaea and eukaryotes, the domain organization of eukaryotic Polε, and the phylogenetic relationship between the other eukaryotic PolBs, both suggest that Polε was a late addition to the nuclear replisome.

Unlike other eukaryotic PolBs, the eukaryotic Polε major subunit is a chimeric fusion of two separate PolB major subunit proteins [4]. The N-terminal portion is catalytically active and homologous to archaeal Polε, while the c-terminal portion is non-catalytic and of unclear origin. The high level of sequence divergence in the non-catalytic c-terminal domain impedes its phylogenetic classification, but structural analysis suggests that it is most closely related to eukaryotic Polα [2]. In yeast, the N-terminal, archaeal-like portion of Polε is nonessential, while the C-terminal portion plays an essential role in replication initiation [5]. This suggests that the non-catalytic domain may have a more ancient origin than the catalytic domain, possibly arising through a duplication of Polα.

In contrast, Polδ enzymatic activity is essential for viability. While it has long been accepted that Polδ is the primary replicator of the lagging strand and Polε is the primary replicator of the leading strand, more recent work suggests that Polδ has a major and essential role in leading strand synthesis [6,7]. One model to reconcile this discrepancy speculates that Polα must hand-off leading strand synthesis to Polδ as an intermediate before Polε can take over leading strand synthesis [6]. Greater integration of Polδ into eukaryotic replisome interactions is also seen by Polδ's reliance on PCNA. Polε's processivity is mostly independent of PCNA, while Polδ lacks processivity without PCNA. Given PCNA's role as an interaction hub of the eukaryotic replisome and as a processivity factor in archaea, less reliance on PCNA suggests a lesser degree of integration of Polε enzymatic activity into the eukaryotic replisome.

In addition to Polε catalytic activity being less integrated into the eukaryotic replisome, the Polε catalytic domain is also a phylogenetic misfit among eukaryotic PolBs. Previous work has shown that all eukaryotic PolBs aside from Polε exist in a large 'Polδ-like' clade [2] and our results broadly recapitulate that organization (Figure 1). The Polε catalytic domain is thus less integrated into the eukaryotic replisome and phylogenetically distinct from other eukaryotic PolBs. Together, these lines of evidence suggest that the archaeal derived portion of Polε was a relatively late addition to the eukaryotic replisome.

Despite being present in close relatives of the eukaryotic cellular ancestor, we suggest that Polε was acquired later through a horizontal acquisition. This scenario would be consistent with the nuclear import step of viral eukaryogenesis (Figure 3B). A nuclear genome, replicated primarily by a viral derived Polδ protein would acquire the Polε catalytic domain from an archaeal cytoplasmic genome, and fusion to a nuclear Polα-like polymerase would redirect Polε activity to the nuclear genome. Acquisition of a second processive polymerase could be particularly advantageous as the nuclear genome grew larger, taking in more cellular genes.

### Viral eukaryogenesis is consistent with accepted evolutionary processes

Although the model of viral eukaryogenesis that we present in Figure 3B may appear to contrast sharply with other models of eukaryogenesis, it is important to stress that the pathway from infectious virus to nucleus is completely consistent with our current understanding of evolutionary processes. In the main text we highlight the phylogenetic evidence for a viral ancestry of several key eukaryotic genes involved in replication and transcription, and we discuss how the compartmentalization of DNA replication and transcription that occurs in both virus factories and eukaryotic nuclei are also consistent with a viral origin of the nucleus. In addition, the linear chromosomes of eukaryotes, which are consistent with those of giant viruses but contrast sharply with the circular chromosomes of archaea, also bear signatures of a viral origin. Moreover, multiploidy is also potentially a viral-derived trait; While the classic image of a provirus is integrated within the host chromosome, recent work has shown that a plasmid lifestyle is common among many bacteriophages. A number of bacteria have multipartite genomes, and their additional chromosomes are thought to have evolved originally from mobile plasmids that acquired essential genes [8]. Acquisition of some essential function by a compartmentalized viral genome would prevent loss of that genome allowing further import of cellular genes to take place. Such import is a known phenomenon, as the majority of mitochondrial DNA has been imported into the nucleus in eukaryotes.

A natural question that arises from the viral eukaryogenesis model is why the cell would evolve towards total compartmentalization of the genome rather than a breakdown of the VF and integration of the viral genome into a cytoplasmic cell chromosome. A possible explanation is that DNA compartmentalization would provide protection against many viruses that replicate within the cytoplasm. Compartmentalization of DNA replication would sequester replication resources away from viruses unable to breach the nucleus, and compartmentalization of transcription would help prevent viral manipulation of host gene expression. Perhaps most importantly, abortive infection and programmed cell death are common defense mechanisms against viruses in both prokaryotes and eukaryotes [9,10]. Compartmentalization of the host DNA sensitizes these systems to the presence of any viral DNA in the cytoplasm, providing potent protection against viral infection. Compartmentalization of DNA replication and transcription has strong implications for viral immunity and may be the driving force for the difference in eukaryotic viromes, which primarily comprise RNA viruses, and prokaryotic viromes, which are dominated by DNA viruses [11].
